# Host population diversity as a driver of viral infection cycle in wild populations of green sulfur bacteria with long standing virus-host interactions

**DOI:** 10.1101/2020.03.05.979559

**Authors:** Maureen Berg, Danielle Goudeau, Charles Olmsted, Katherine D McMahon, Jennifer Thweatt, Donald Bryant, Emiley A Eloe-Fadrosh, Rex R Malmstrom, Simon Roux

## Abstract

Viral infections of bacterial hosts range from highly lytic to lysogenic, where highly lytic viruses undergo viral replication and immediately lyse their hosts, and lysogenic viruses have a latency period before replication and host lysis. While both types of infections are routinely observed in the environment, the ecological and evolutionary processes that regulate these different viral dynamics are still not well understood. In this study, we identify and characterize the long-term dynamics of uncultivated viruses infecting green sulfur bacteria (GSB) in a model freshwater lake sampled from 2005-2018. Because of the additional requirements for the laboratory cultivation of anaerobes like GSB, viruses infecting GSB have yet to be formally identified, leaving their diversity and impact on natural populations of GSB virtually unknown. In this study, we used two approaches to identify viruses infecting GSB; one *in vitro* based on flow cytometry cell sorting, the other *in silico* based on CRISPR spacer sequences. We then took advantage of existing bulk metagenomes derived from Trout Bog Lake covering the 2005-2018 period to examine the interactions between GSB hosts and their viruses across multiple years and seasons. From our data, GSB populations in Trout Bog Lake were found to be concurrently infected with at least 2-8 viruses each, many of which were lysogenic viruses; one GSB host population in particular was consistently associated with two lysogens with a nearly 100% infection rate for over 10 years. We illustrate with a theoretical infection model that such an interaction can be stable over multiple years given a low, but persistent level of lysogen induction in host populations with already high infection rates. Overall, our data suggest that single GSB populations are typically infected by multiple viruses at the same time, that lytic and lysogenic viruses can readily co-infect the same host population in the same ecosystem, and that host strain-level diversity might be an important factor controlling the lytic/lysogeny switch.

## INTRODUCTION

Recent advances in metagenome sequencing have enabled high-throughput exploration of the virosphere, leading to a >200-fold increase in viral genomes available in databases, and uncovering >1,000 new genus-level viral groups (Emerson et al., 2018; Paez-Espino et al., 2016; Roux et al., 2016; Suttle, 2007). While these data provide new insights into viral genome diversity,most of these uncultivated viral genomes are not associated with any host. The scarcity of host association for uncultivated viruses means that basic viral ecology parameters, such as how many different viruses infect a single host population and how the diversity of virus and host populations change over time, are poorly constrained. Correspondingly, this limits our ability to fully understand virus-host interactions in nature, and severely hinders efforts to integrate viruses into ecosystem models.

Viral infection dynamics exist along a spectrum from highly lytic to lysogenic and/or chronic (Weitz et al., 2019); highly lytic viruses undergo viral replication and host lysis immediately, while lysogenic viruses have a latency period before replication and host lysis. However, the ecological and evolutionary mechanisms affecting these different infection dynamics are still unclear. Specifically, one of the most fundamental questions in viral ecology that remains unanswered is which environments and/or life-history traits select for more lytic or lysogenic viruses. This is an actively researched and discussed topic, both in a viral ecology framework (Knowles and Rohwer, 2017; Knowles et al., 2016; Silveira and Rohwer, 2016; Weitz et al., 2017; Winter et al., 2010), and a molecular biology framework (Erez et al., 2017; Silpe and Bassler, 2019). Building from a combination of experimental virus-host systems and whole-community studies, different hypotheses about these ecological and evolutionary drivers have been proposed. The “kill the winner” model places importance on the lytic life stage of viruses, and proposes that when a host is actively growing, host population numbers increase, and lytic viruses will be favored over lysogenic viruses (Thingstad, 2000). In this scenario, lytic viruses suppress host blooms, and maintain diversity in the host population by preventing one host strain from outcompeting all others (Knowles et al., 2016). Lysogeny is seen as an “alternative” choice in this model, for when hosts are not as readily available. Others have proposed similar views by postulating that lysogeny is governed by host trophic status. For example, lysogeny seems to be favored in some environments when host productivity is low, or for hosts with variable growth rates, such as pathogenic microbes or environments with strong seasonal patterns (Brum et al., 2016; Payet and Suttle, 2013; Touchon et al., 2016).

In contrast, the “piggyback-the-winner” model proposes that lysogeny is favored when hosts are readily available. This may be advantageous in environments with fast growing hosts, where the viruses would profit more from lysogeny than from host lysis, for example. This scenario reduces the amount of control viruses could exert over bacterial abundance, and places more importance on other types of host interactions, e.g. superinfection exclusion conferred by the lysogens (Knowles et al., 2016). The existence of such diverging hypotheses, both with supporting data, reflects the complexity of virus-host interactions and the limits of our current understanding. In particular, the majority of the hypotheses put forward so far have been based on community-wide measurements, which average out differences associated with individual viruses and host populations. In addition, most focus on ecophysiological traits, such as host growth rate and nutrient availability, and only a few consider host and virus population diversity, although virus-host coevolution at the strain level have been described (Koskella and Brockhurst, 2014; Poullain et al., 2008; Rodriguez-Valera et al., 2009). In order to expand our understanding of these host-virus dynamics, high-resolution datasets that incorporate a variety of specific host-virus pairs across time and environments are required.

In this study, we identify and characterize the long-term dynamics of uncultivated viruses infecting green sulfur bacteria (GSB) in a model freshwater lake (Trout Bog Lake) sampled from 2005-2018. GSB are anoxygenic photoautotrophic bacteria that can form massive blooms in illuminated, low oxygen/high sulfide waters, and they play a central role in global carbon and sulfur cycling (Frigaard and Dahl, 2009; Gregersen et al., 2009; Vogl et al., 2012). Viruses likely exert a strong influence on aquatic GSB populations by controlling bloom dynamics and through horizontal gene transfer (HGT), especially since comparative genomics previously revealed an extensive history of HGT across GSB (Frigaard and Bryant, 2008; Llorens–Marès et al., 2017). GSB have additional requirements for anaerobic cultivation and growth in the laboratory, and therefore viruses infecting GSB have yet to be formally identified, leaving their diversity and impact on natural populations of GSB still relatively unknown. In this study, we used two approaches to identify viruses infecting GSB. The first uses *in vitro* based on flow cytometry cell sorting, the other *in silico* based on CRISPR spacer sequences. We next leveraged existing bulk metagenomes of Trout Bog Lake covering the 2005-2018 period to examine the interactions between GSB hosts and their viruses across multiple years and seasons. Our data suggest that single GSB populations are typically infected by multiple viruses at the same time, that lytic and lysogenic viruses can readily co-infect the same host population in the same ecosystem, and that host strain-level diversity might be an important driver of the lytic/lysogenic switch.

## MATERIALS AND METHODS

### DNA sampling and sequencing

Trout Bog Lake is located in Wisconsin, USA, and is surrounded by a sphagnum mat that supplies large amounts of terrestrially derived organic matter to the lake, leading to darkly stained water and greatly attenuated light penetration. This dimictic lake has a maximum depth of 9 m, surface area of ~11,000m^2^, and a mean pH of 5.1. Dissolved oxygen is below detection in the hypolimnion between ice thaw (early May) and fall mix (mid-November). Two different types of samples were collected during June-October 2018: (1) depth-integrated water samples were collected from the hypolimnion layer (as defined by the temperature gradient) at 14 different time points, and (2) depth-discrete water samples were collected at every meter throughout the water column at 14 different time points. Both types of samples were filtered onto a 0.2μm pore-sized polyethersulfone filters (Supor 200, Pall, Port Washington, NY) prior to storage at −80°C. DNA was purified from these filters using the FastDNA Spin kit for Soil (MP Biomedicals, Santa Ana, CA), with minor modifications (Shade et al., 2007). For the depth-discrete water samples, a subsample of water was preserved using glyTE and analyzed through flow cytometry (see below).

For the depth-integrated samples, one library from each sample was prepared following standard Illumina TruSeq protocol. For the depth-discrete samples, Nextera XT v2 (Illumina) sequencing libraries were generated from 5,000 sorted cells following multiple displacement amplification (MDA). Five libraries of sorted Green sulfur bacteria, i.e. pigmented cells that were stained by SYBR Green (Invitrogen) (488nm excitation; 530nm emission) and emitted autofluorescence >750nm after excitation with 642nm laser (Figure S1), and two additional libraries of sorted nonpigmented cells were sequenced for each sample. Sorted cells were amended with glycerol (6% final conc.) and stored at −80°C while awaiting MDA. Prior to amplification, cells were pelleted by centrifugation at 6,350 x g for 1h at 10°C, inverted and spun at 7 x g for 5s to remove supernatant, and amplified by MDA at 45°C for 16hrs using EquiPhi29 DNA polymerase (ThermoFisher Scientific) following the protocol described in Stepanauskas et al., 2017). Paired-end sequences of 2 x 150 bp were generated for all libraries on the NovaSeq platform (Illumina). Metagenomic sequence reads are publicly available on the JGI IMG portal.

### Data processing and genome binning

For depth-discrete water samples, each metagenome was subsampled to 60 million total reads per sample using BBMap reformat function to generate our “targeted metagenomes” (v37.92, Bushnell, 2014). For each sampling date, reads from the 5 replicates were pooled and coassembled using SPAdes (v3.10.1, Bankevich et al., 2012). For each coassembly, both positive and negative cell fractions were read-mapped to the contigs using Bowtie2 (default parameters, (v2.3.4.3, Langmead and Salzberg, 2012)). To remove any cross-talk contamination during sequencing, only contigs that had at least 10x higher coverage in the positive fraction (compared to the negative fraction) were kept for downstream analysis. For each coassembly, after filtering, contigs were binned based on CONCOCT clustering using Anvi’o (v5.4, Eren et al., 2015) and bin quality was determined using CheckM (v1.0.12, Parks et al., 2015). Once binning was complete for each coassembly, all bins were pooled and dereplicated using dRep (v2.2.3, Olm et al., 2017). The dereplicated genome set was used as the representative genomes across all sampling dates in 2018. Both depth-integrated and depth-discrete were read-mapped to these representative metagenomes using Bowtie2, and coverage data was extracted using Anvi’o.

### Time series dynamics

Depth-integrated samples (referred to as “bulk metagenomes” in this text) were previously taken from Trout Bog Lake in 2005, 2007-2009, and 2012-2013 (Bendall et al., 2016). In order to conduct a time series analysis of Green sulfur bacteria and their viruses, reads from those samples were read mapped to both the representative GSB genome set and the representative GSB virus set (see below) using Bowtie2. Coverage data was extracted using Anvi’o.

### Phage identification and characterization

To identify potential viral contigs, contigs from all 14 co-assemblies of sorted cells and all bulk metagenome assemblies were analyzed by VirSorter 1.0.5 (Roux et al., 2015a). Briefly, VirSorter looks for hallmark viral genes (e.g., tail proteins, terminase, capsid protein) and enrichment of viral-like genes in the contigs provided, and ranks the contigs based on these characteristics along with matches to known viral genomes. Sequences identified by VirSorter in categories 1, 2, 4, and 5, were manually curated, and genuine viral contigs from each coassembly were pooled and dereplicated to produce a representative set of potential green sulfur bacteria viruses.

To identify contigs from putative GSB viruses using matched CRISPR spacers, all metagenomes were first analyzed with the Crass assembler (v0.3.12, Skennerton et al., 2013) to assemble CRISPR arrays. Then, spacers associated with repeats identical to the ones detected in GSB genomes (Table S1) were gathered and compared to all viral contigs identified across all metagenomes (see above) using blastn with options optimized for short sequences (“-dust no - word_size 7”). Predicted viral contigs showing at least one match to a GSB-associated spacer (0 or 1 mismatch across the whole spacer length) were selected as “candidate GSB viruses”.

Both depth-integrated and depth-discrete libraries were read-mapped to these viral contigs using Bowtie2, and coverage data was extracted using Anvi’o. For taxonomic affiliation, all viral contigs identified (including the ones not associated with GSB) with a length ≥10kb were pooled with reference genomes from NCBI RefSeq (v93, filtered to include only genomes from bacterial and archaeal viruses), along with viral sequences from the IMG/VR database with a predicted GSB host (Paez-Espino et al., 2019), prophages predicted in GSB genomes from the IMG database (REF, sequences predicted with VirSorter, categories 1/2/4/5), and viral sequences ≥10kb identified in metagenomes from Lake Mendota (predicted with VirSorter as for Trout Bog Lake metagenomes, see above). The additional datasets beyond RefSeq were included to try to ensure a large representation of sequences from freshwater viruses and/or putatively associated with GSB. This set of viral genomes and contigs was first dereplicated (95% average nucleotide identity [ANI], 85% alignment fraction) using MUMMER 4.0.0b2 (Marçais et al., 2018). The resulting dataset was then used as input to build a genome network based on shared gene content using vContact 2 ((Bolduc et al., 2017), using diamond for all-vs-all protein comparison, MCL for protein clustering, and ClusterOne for genome clustering). The resulting network was visualized using Cytoscape (Shannon et al., 2003).

### SNP identification and analysis

SNPs were identified using Anvi’o. Only SNPs with at least 10x coverage were retained for analysis. Allele frequencies were rarified to a depth of 10, and SNPs were filtered again to exclude those SNPs whose dominant allele frequencies were now 1. Nucleotide diversity (Pi) was calculated for each SNP, values were summed (per sample), and divided by genome size. SNP turnover was calculated in a similar way, although specific years were compared to each other, rather than all together.

### Host-virus model

The model was based on Weitz et al., 2019, with *p* (prophage decay rate, h^-1^) and *cost* (growth cost) as additional parameters, and includes the following nonlinear ordinary differential equations:

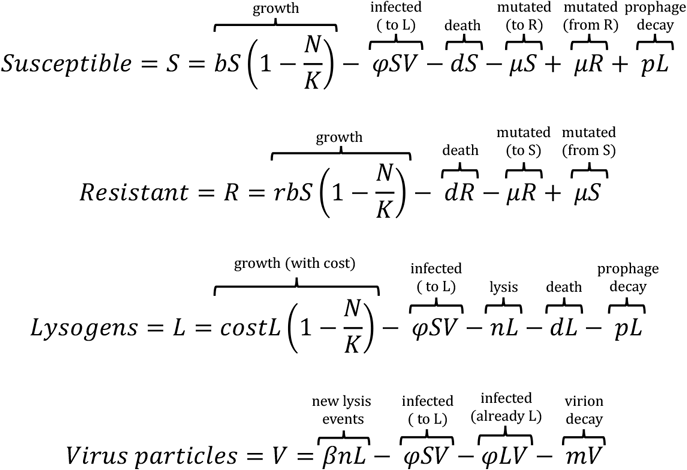

S, R, L, and V denote the densities of susceptible cells, resistant cells, lysogenic cells, and virus particles per ml, respectively. N denotes the total density of cells, i.e. S + R + L. The parameters similar to Weitz et al. 2019 include *b* (maximal cellular growth rate, h^-1^), *K* (carrying capacity, ml^-1^), *φ* (adsorption rate, ml/h), *d* (cellular death rate, h^-1^), *β* (burst size, number of virus particles), *m* (virion decay rate, h^-1^), and *n* (induction rate, h^-1^). Additional parameters are *μ*R (mutation rate from susceptible to resistant, h^-1^), *μ*S (mutation rate from resistant to susceptible, h^-1^), *p* (prophage decay rate, h^-1^), *r* (effect of resistance on cell growth), and *cost* (effect of lysogeny on cell growth). Cell death (unrelated to phage infection) was kept identical across all cell types. The adsorption of virus particles to susceptible and lysogenic cells was also kept identical, with the latter being an evolutionary dead-end (i.e. no new lysogenic cells or virus particles are produced from these events), and the adsorption to resistant cell was considered as null (i.e. the resistance was assumed to arise from an absence of adsorption of the virus particle to the resistant host cell).

The two parameters which varied between computations were *n* (induction rate) from 1×10^-6^ to 1 and the initial proportion of lysogenic cell from 0 to 100%. The initial number of cells was set at 1×10^5^ ml^-1^ based on the flow cytometry counts of GSB for the first samples in our time series, while the maximum carrying capacity K was set at 3×10^-6^ based on the maximum count of GSB throughout the time series (see above). The initial number of viral particles was set at 1×10^2^ ml^-1^. Other parameters were as follows: maximum growth and death rates were 0.1 and 0.024, respectively, which allows for a bloom of GSB to form in days when not including any virus particle or lysogenic cell. Burst size, virion decay, and induction rate were set as in Weitz et al., 2019, i.e. *β* = 50, *φ* = 6.7×10^-10^ ml/h, and *m* = 1/24 h^-1^. Prophage decay rate *p* was set at 1×10^-4^ h^-1^ corresponding to ~1% of prophage loss per generation at the maximum growth rate of 0.1 h^-1^, in the same order of magnitude as the rate of prophage loss observed for some laboratory systems (Alexeeva et al., 2018). Mutation rate from susceptible to resistant (*μ*R) and from resistant to susceptible (*μ*S) was arbitrarily set at 1×10^-6^ to ensure a constant pool of both susceptible and resistant host cells in the system. Finally, the adjusting parameter for lysogenic growth rate was set at 0.99 to reflect the added energetic cost of the integrated prophage, and the adjusting parameter for resistant cells growth rate was set at 0.90 to allow for the establishment of typical prey-predator dynamics for high induction rate (i.e. lytic phages).

## RESULTS

### Green sulfur bacteria diversity in Trout Bog Lake

During the summer and early fall of 2018, Trout Bog Lake was sampled on a nearly weekly basis to generate the metagenomic data introduced in this paper. Two types of sampling were performed: one targeted green sulfur bacteria (GSB) by sampling specific depths of the water column (depth-discrete), and the other aimed to collect the entire microbial community by sampling large segments of the water column (standard bulk metagenomes, see Methods). Using the depth-discrete samples, the relative abundance of GSB was quantified using FACS across all sampling dates in 2018 (Figure 1A); while other bacteria have this type of fluorescence, our sequencing results (below) suggest that GSB were the only bacteria that were sorted, supporting our use of FACS data as a measurement for GSB abundance. Early in the summer, we measured ~140,000 GSB cells/mL, and GSB peaked in mid-August with 2.2 million GSB cells/mL (32% of total cells screened). The overwhelming majority of GSB cells were found below the oxygen barrier/chemocline (white line in Figure 1A), consistent with the anaerobic metabolism of GSB. This type of seasonal pattern for GSB has been observed in another freshwater lake systems, including through 16S rRNA gene amplicon sequencing data (Diao et al., 2018), where GSB were dominant during the summer when the lake developed stratified layers. For each of our sampling timepoints (using depth-discrete samples), we generated 5 replicates of GSB-targeted metagenomes (5,000 cells sorted based on size and autofluorescence, see Figure S1), along with 2 replicates of non-GSB metagenomes (5,000 cells positive for DNA stain, but negative for autofluorescence), for each time point and from the depth at which the GSB cell count was highest. Similar datasets were also generated for one sampling date in Fall 2017 (see Methods).

**Figure 1.**
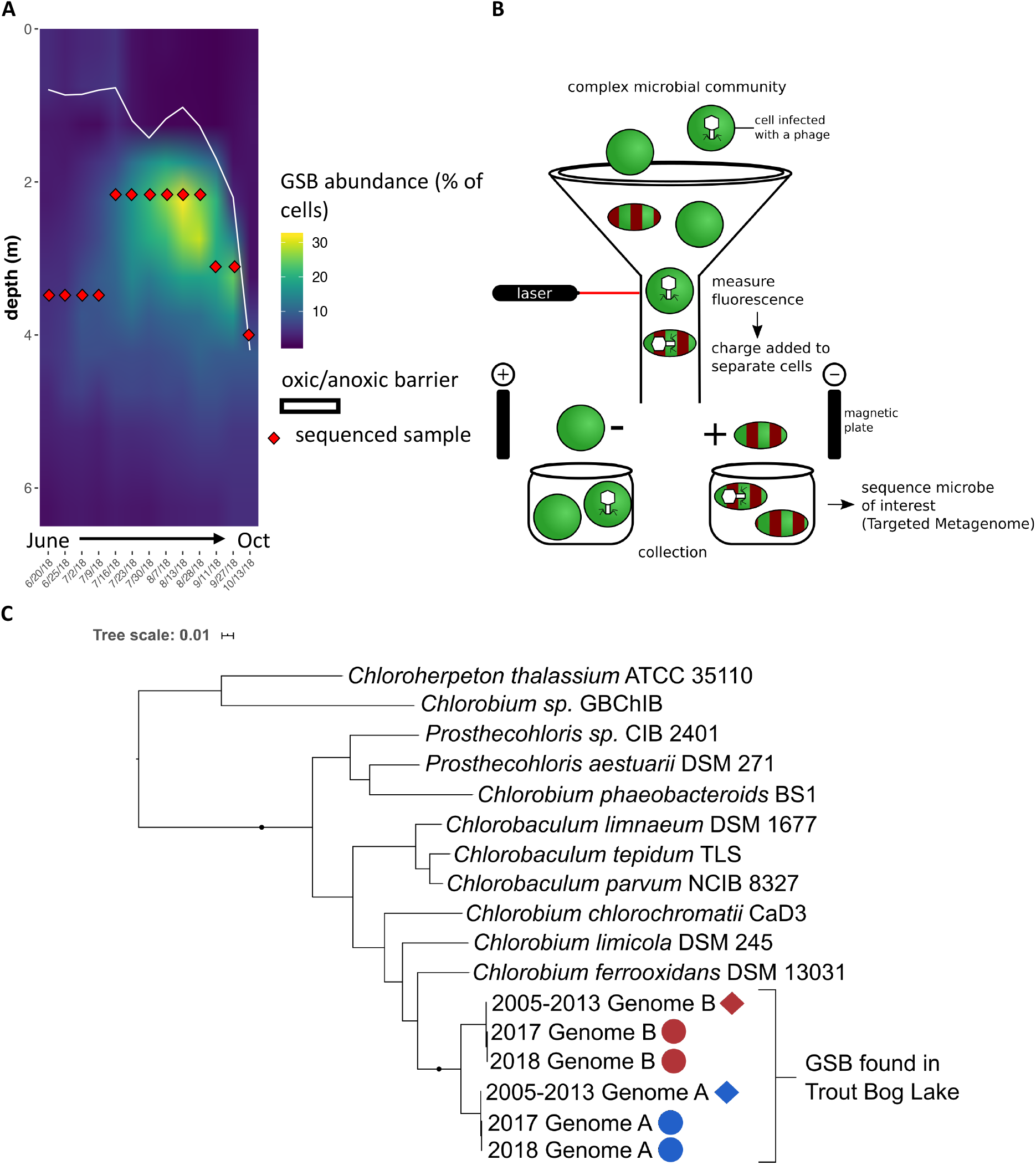
**A)** An average of 3.7 million cells/mL were measured for per sample per date; shown is GSB abundance (% of total cells per sample per date). Red diamonds represent the specific depth that was sequenced. The white line represents the oxic/anoxic barrier. **B)** GSB cells were sorted using FACS, targeting green DNA stain and red autofluorescence. **C)** Phylogenetic tree based on rpoB genes of GSB from Trout Bog Lake and GSB isolate genomes; circles represent new genome bins from our targeted metagenomics, and diamonds represent previously published genome bins; colors represent Genome B (red) and Genome A (blue).

Bacterial genomes were binned from a combined assembly of GSB-targeted metagenomes from each sampling date, resulting in 14 sets of genome bins (one set per date). Each sampling date included 1 or 2 nearly complete GSB genome bins (completeness estimation: 97-100%, redundancy estimate: 0-2%, Table S2), with no other taxa binned in our sorted samples. Based on Average Nucleotide Identity (ANI) clustering, bins from all 14 sampling dates were found to represent 2 distinct genomes, hereafter designated as “GSB-A” and “GSB-B” (Table S3A). Previous exploration of GSB community diversity in lakes using methods such as amplicon sequencing and fingerprinting also typically found only a few different GSB in each location,suggesting that only a few dominant GSB populations occur in a given environment (Diao et al., 2018; Martínez-Alonso et al., 2005; Vila et al., 2002). When placed in a phylogeny with known isolate genomes and previously published Trout Bog Lake genome bins, both GSB-A and GSB-B branched within the *Chlorobiaceae* family, and tightly clustered with the previously described Trout Bog Lake GSB genomes (Figure 1C) (Bendall et al., 2016). ANI comparison confirmed that both GSB genome bins recovered from targeted metagenomes were the same populations as those found in the bulk metagenomes throughout 2005 to 2013 (ANI > 99.5%). Notably, GSB-A corresponds to a GSB population that was previously observed to undergo a genome-wide sweep between 2005 and 2009 (Chlorobium-111 in Bendall et al., 2016, Table S3B,C).

Because we targeted the GSB maximum depth at each time point (to sample the depth with maximum GSB concentration), targeted metagenomes were sequenced from different depths across 2018, highlighting differential distribution patterns for GSB-A and GSB-B (see Figure 1A). Specifically, GSB-A was only recovered at lower depths (3-4m), while GSB-B was recovered at all sequenced depths, including higher in the water column during the summer bloom (2.5-4m, Figure S2, Table S2). This suggests that the co-existence of these two GSB populations in Trout Bog Lake is possibly linked to niche partitioning along the water column. This would be consistent with the genomic differences between GSB-A and GSB-B, which despite being classified as two closely related species, display ~23% unique gene content with respect to each other. This type of niche partitioning has been seen in GSB before, with brown-pigmented GSB blooming deeper than green-pigmented GSB (Crowe et al., 2017; Montesinos et al., 1983); this type of laying can also be found within green-pigmented GSB, depending on which specific pigments they produce (Maresca et al., 2004). Based on pigment gene analysis, both our GSB populations appear to be green-pigmented GSB. It is unclear if they produce the same green pigments, as the same set of pigment genes can be found in both genomes, although the actual sequences of these genes are not identical between the two GSB.

A third bin was routinely found in the 2018 targeted metagenomes, however it was estimated to be only 20% complete, i.e. it includes very few to none of the expected core bacterial genes. This third bin was clustered next to the other two GSB bins when included into a phylogeny, suggesting it also represents a GSB population. A similar genome was also binned in the 2017 targeted metagenomes, but never in the bulk metagenomes from 2005-2013 even though read mapping revealed that the corresponding sequences were present in those datasets at a low relative abundance, suggesting that this third bin could represent a distinct and rare GSB population. Considering that we never recovered a complete genome the third bin, and that no viruses were found in this bin, only GSB-A, GSB-B, and their associated viruses were further analyzed.

### GSB viruses recovered from targeted metagenomes

To identify viruses infecting GSB, we first used VirSorter to detect viral contigs from the 2017 and 2018 GSB targeted metagenomes, and excluded putative contaminants based on coverage patterns across replicates (see Methods). Across all 14 sampling dates, we recovered 43 non-redundant predicted GSB viral contigs (clustered at 95% ANI – 80% Alignment Fraction, AF) (Table S4). While most of these viral sequences were short and could represent decayed prophages or rare viruses that might be difficult to assemble, we identified 11 complete or nearly complete GSB virus genomes, on which we focused our analysis. Out of the 11 viruses, 10 were identified as temperate (i.e. able to enter a lysogenic cycle) because they encoded an integrase gene and/or were assembled as an integrated prophage within our GSB genomes. The other viral contig, CV-1-33, did not contain an integrase gene, nor was it ever assembled as integrated into the host genome, and thus it is probably not a temperate virus. Only one putative GSB viral sequence was described in the past, identified as a virus infecting GSB based on metagenomic coverage, however it bears no similarity to the GSB viruses identified here (Llorens–Marès et al., 2017). Putative GSB viruses were also recruited from all available Trout Bog Lake metagenomes from 2005-2018 by matching spacers from the CRISPR arrays associated with GSB-A and GSB-B (see Methods). This yielded a total of 534 contigs, which were subsequently clustered into viral OTUs (95% ANI and 85% AF) to produce 45 non-redundant viral contigs (34 of the contigs were > 10kb). Out of the 45 contigs, 6 matched viruses recovered through flow sorting GSB. Overall, combining results from the flow sorting method and the CRISPR spacer-matching method resulted in 50 non-redundant GSB viruses.

To situate these new GSB viruses within the global virus diversity, we used a genome-based network analysis of their shared protein content with vContact 2. The majority of the new GSB viruses (45/50) were connected to the main component of the network, confirming that these are likely members of the *Caudovirales* order (Figure 2, Figure S3). However, they formed novel clusters (approximately genus/subfamily rank), which did not include any other reference sequences beyond GSB viruses. This is consistent with the lack of isolated GSB viruses, and the fact that viruses tend to cluster by large host groups (~phylum/class) in gene-content-based networks (Roux et al., 2015b). Within the GSB viruses, most sequences tend to cluster by themselves (as singletons), with two exceptions where GSB viruses formed their own clusters. These two clusters of GSB viruses showed different patterns. In sub-cluster VC-2, three genomes assembled from the 2018 targeted metagenomes were clustered with a 34-59% shared gene content; of the three, one was binned with and co-detected with GSB-A, while the two others were binned and co-detected with GSB-B. Based on this shared gene content, the GSB viruses in this sub-cluster appear to be recently diverged and co-diversifying each with a specific host population (Figure 2). Conversely, in sub-cluster VC-1, one genome assembled from the 2018 targeted metagenomes was clustered with one genome from the 2017 targeted metagenomes and one from the 2007 bulk metagenomes (Figure 2). Comparisons of these three variants revealed two distinct regions of the virus genome: a conserved region with structural genes such as capsid and tail genes, and a variable region with mostly uncharacterized proteins. Hence, these three distinct genomes likely represent different variants from the same population with different relative abundance through the years. This pattern of highly variable regions across years is an exception among GSB viruses: for the other 3 viruses identified in the targeted metagenomes discussed in this paper, identical or near-identical sequences (>95% ANI over > 85% AF) could be identified in the 2005-2013 bulk metagenomes (Figure 3, Figure S4).

**Figure 2.**
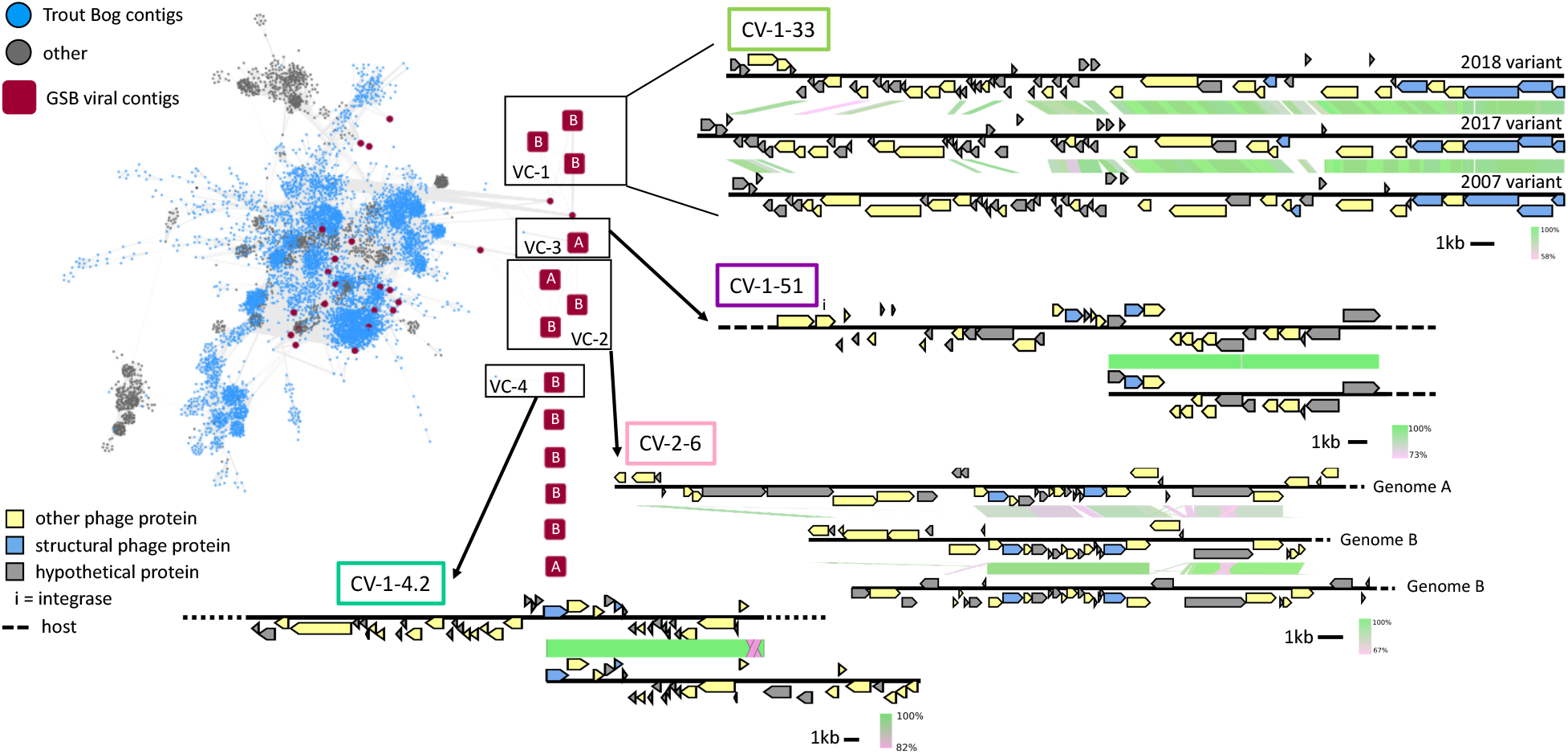
Putative viral contigs (blue circles; red squares GSB-specific viral contigs) from Trout Bog Lake (TBL) were clustered with viral databases (grey circles). Network edges represent shared gene content between viral contigs. GSB viral contigs clustered into 3 groups (VC −1, −2, −3, and −4). VC-1 contains the three CV-1-33 variants; VC-3 and VC-4 contain CV-1-51 and CV-1-4.2, respectively, plus a previously sequenced TBL metagenome viral contig. Genome alignments were generated using blastn, with green representing 100% nucleotide alignment. Gene content is color-coded, and dashed lines show regions where the host genome was found. All GSB viral contigs are labeled with their host (either “A” or GSB-A or “B” or GSB-B).

**Figure 3.**
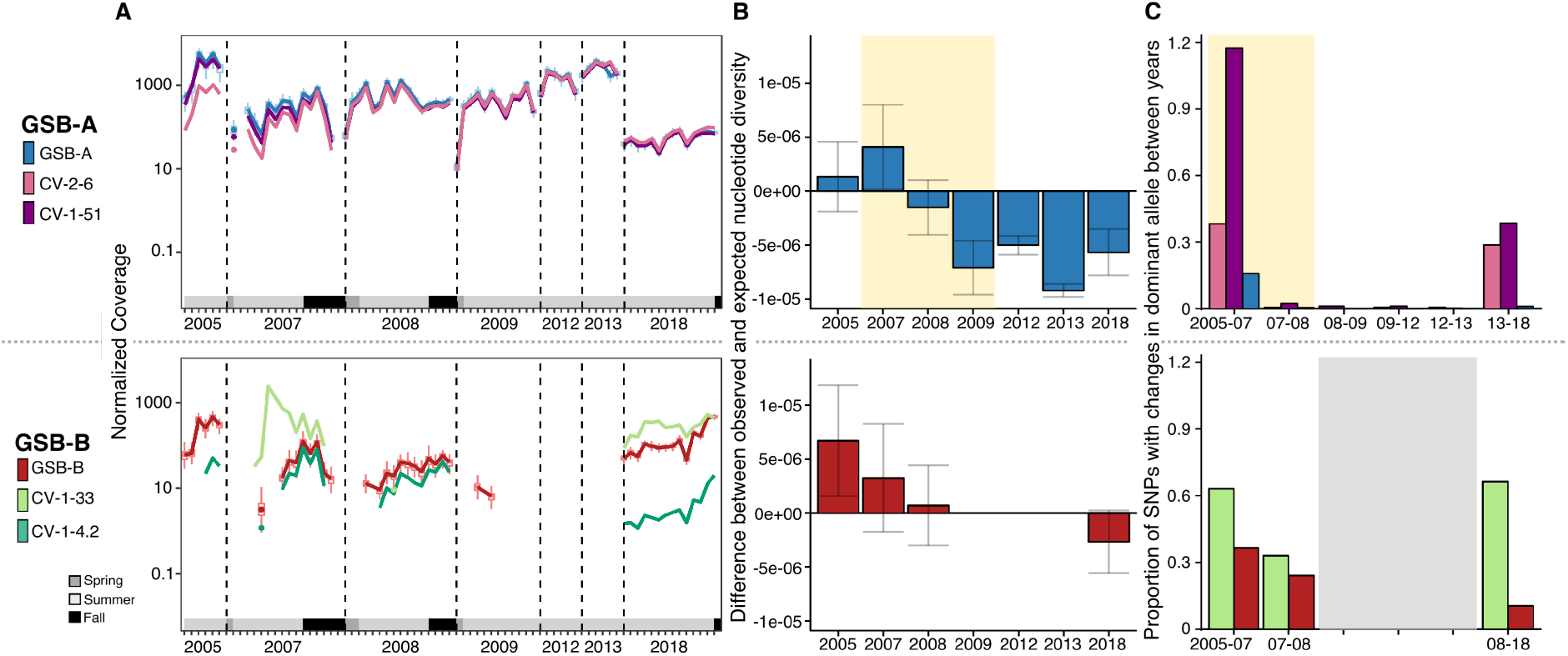
**(A)** lysis rate against ratio of lysogenized host cells,showing infection rate (top) or resistant host density (center); color scales on the right for each respective plot; (bottom) shows a summary schematic for the above plots. CV-1-51 and CV-1-33 are labeled to represent the conditions that generated the vignettes **(B)** Difference between observed and expected nucleotide diversity for GSB-A and GSB-B; expected values are calculated using a regression of observed nucleotide diversity and raw (not normalized) coverage; bars show standard deviation; yellow shaded region denotes time period during genome sweep. **(C)** Proportion of SNPs that change between years of GSB-A and associated viruses and of GSB-B and associated viruses. For diversity and SNP assessments, years were excluded from analysis if there was less than a 10x coverage for each SNP; yellow shaded region denotes time period during genome sweep.

Overall, our identified GSB viruses are distinct from all known viruses (including one putative GSB virus from Llorens–Marès et al., 2017)), may represent six putative new genera, and appear to be genetically stable across decades except for one population, which showed large changes within the same time frame in a hypervariable region. This high degree of genetic stability across decades was unexpected, as most work on viral genome evolution has shown rapid genome change (of varying degrees) across time, similar to what was seen in our VC-1 (which comprises virus CV-1-33) (Ignacio-Espinoza et al., 2019; Mavrich and Hatfull, 2017; Paez-Espino et al., 2015). It is possible that these genetically stable GSB viruses are no longer able to be induced and enter into the lytic cycle, and have become a part of the host genome, but in those instances, we would expect to see degradation of the prophage instead of high stability across multiple years (Ramisetty and Sudhakari, 2019).

Most (26) of the 45 viral OTUs detected in 2005-2013 based on CRISPR matches were never (or rarely ever) detected in 2018, either in targeted or bulk metagenomes (Figure S4). A temporal comparison of viral contig coverage and the acquisition of their respective spacers revealed multiple instances of decline in virus coverage following spacer acquisition, suggesting successful defense by the host population against these infections (Figure S4). Interestingly, we found a variety of CRISPR-virus dynamics within our relatively simple host-virus system: viruses with no spacers (CV-1-51), viruses with unique spacers recovered every year (CV-2-6), viruses present for many years before spacers can be detected (mg13_3-398034), viruses that are found long after their spacers are no longer detectable (mg05_20-709165), spacers that are retained even after the virus seemingly disappears (mg18_1120), and viruses that disappear after the presence of spacers (mg09_208) (Figure S4B). The diversity of CRISPR-virus dynamics in this one system suggest that spacers may have different levels of efficacy within a single host, different frequency within a host population, and that eventually, individual virus-host pairs can be associated with a broad range of CRISPR dynamics.

The fact that different GSB viruses are identified through targeted metagenomes and CRISPR spacer matching highlights how these two approaches are complementary, as they reflect different infection status. While CRISPR-based recruitment identified many putative GSB viruses, most did not seem to represent ongoing, ecologically relevant infections but instead reflected past infections. Conversely, viral sequences identified in targeted, flow-sorted metagenomes represent “in-cell” viruses, including integrated prophages not targeted by CRISPR-Cas systems, although this method may be less suitable for very rare (i.e. low rate of infected cells) or transient (i.e. short in-cell time) infections. Together these two approaches identified dozens of viruses infecting GSB over a 13-year time period, and revealed these GSB populations were often simultaneously infected with >15 viruses at a single time point (Figure S4).

Auxiliary metabolic genes (AMGs) are host genes found in bacterial viruses that can modulate host cell metabolism during infection, usually to increase the efficiency of viral replication. This has been demonstrated in many bacteria, including cyanobacteria, where viral-encoded proteins are functional during host photosynthesis (Lindell et al., 2005). In GSB, comparative genomics previously revealed an extensive history of HGT, with the *sox* cluster for thiosulfate utilization as a well-known example (Frigaard and Bryant, 2008), so it is possible that GSB viruses carry AMGs. However, we found no evidence for AMGs in our GSB viral genomes. Because the *sox* gene cluster is a well-known example of HGT, we examined the two GSB populations in this study for thiosulfate utilization capacity. While we see these genes in the community (i.e. metagenomes), we don’t see a definitive conclusion through genome binning that either GSB population consistently or systematically encodes this gene cluster.

### Host-virus temporal dynamics in Trout Bog Lake

Mapping reads from individual bulk metagenomes to the new GSB genomes and GSB viruses assembled from targeted metagenomes enabled a more detailed investigation of the temporal dynamics of GSB and their viruses in Trout Bog Lake. These data provide a unique look at virus-host dynamics, as these data cover more than 10 years, and represent one of the longest running datasets currently available to investigate viral-host dynamics. GSB-A was present in all years dating back to 2005, and was, until 2018, the dominant host population. Its associated viruses (CV-2-6 and CV-1-51, both integrated prophages) appear to be present at a low relative abundance in 2005, and became pervasive in 2007, seemingly infecting every member of the host population from this date on (Figure 3A). While both host and viruses decreased in abundance in 2018 relative to previous years, they were still abundant as judged by coverage (>40x; ~32% of GSB reads in 2018 were GSB-A) and showed the expected increase in summer compared to spring samples corresponding to the GSB bloom. While this long-term virus-host stability was unexpected, as viruses are often thought of as fast-evolving, recent work done in marine systems also showed long-term stability in viral communities, resulting in long-term virushost coexistence (Ignacio-Espinoza et al., 2019). The mechanisms for this stability are likely different however, as those coastal marine systems seem to rely on perpetually changing minor variants (Red Queen Hypothesis, Brockhurst et al., 2014), while no evidence of such an arms race was observed in Trout Bog Lake GSB-A viruses.

In contrast, GSB-B was overall less abundant, with reliable coverage (>10x) only achieved in 2005, 2018, and some samples in 2007. While it was not the dominant GSB population for most of the years covered by our data, it was clearly more abundant than GSB-A in 2018 (Figure 3A).

Similar to GSB-A, GSB-B is also associated with an integrated prophage (CV-1-4.2). However, unlike the prophages that infect GSB-A, CV-1-4.2 was quite rare, likely infecting only a subset of the population (Figure 3A). The other GSB-B virus (CV-1-33) displayed a strikingly different virushost dynamic than all the other GSB viruses we observed, with large peaks in abundance compared to its host in 2007 and 2018, likely the result of ongoing lytic infections. Further, CV-1-33 was routinely and consistently found in the GSB cells, as it was present in 10 out of the 14 sampling dates in 2018 (depth-discrete samples), suggesting active infection. It is notable that CV-1-33 is also the only virus that shows high mutation/recombination rates with a clear hypervariable region (see above and Figure 2), which would be consistent with a lytic virus engaged in an “arms race” with its host. These data also confirm that virus-host association based on relative abundance correlation could be efficient for prophages but may misinterpret the dynamics of lytic viruses (Figure 3), as predicted by theoretical models (Coenen and Weitz, 2018).

### Microdiversity of GSB and associated viral populations

As previously reported in Bendall et al., 2016, GSB-A went through a genome wide sweep in 2007, leading to a stark reduction in population diversity (Figure 3B). The same pattern could be observed here for the associated viruses, which both underwent a clear genome-wide sweep (Figure S5). In 2018, we are able to see some resurgence of population diversity in terms of single nucleotide variants in both the host and the viruses (Figure 3B, Figure S5), suggestive of an ongoing slow diversification process. We were only able to reliably ascertain population diversity in 2005, 2018, and some of 2007 because of the low GSB-B coverage in most years (Figure 3B). However, GSB-B did not seem to undergo a similar genome sweep (Bendall et al., 2016). Viral infections could have been a possible cause of the genome sweep seen in GSB-A, because we can see an increase in prophage abundance concomitant with a decrease in GSB-A population diversity (Figure S6). On the other hand, the genome sweep may have provided the appropriate conditions for the GSB-A viruses to attain high levels of abundance if these two viruses happen to infect the strain of GSB-A that became dominant after the sweep, reminiscent of the “piggyback-the-winner” hypothesis (Knowles et al., 2016).

In addition to estimating microdiversity within GSB hosts and their associated viral populations each year, we also looked at the turnover between populations each year by calculating the proportion of SNPs that changed between years. This was calculated similar to our standard SNP calculations, but instead of looking at overall SNPs density across all samples, pairwise comparison of years was computed. By looking at turnover in the GSB populations, we can estimate how similar (or dissimilar) a given population is compared to the population in the year before or after. In GSB-A, we see a similar pattern as the within-sample nucleotide diversity, with higher SNP turnover between 2005 and 2007, and also between 2013 and 2018 (Figure 3C), suggesting that the GSB populations within these time periods were less similar than between 2007 and 2013. Between 2007 and 2013, the same clonal GSB-A population dominated every year, as little-to-no SNP changes between or within years were observed. For the GSB-A associated viruses, we see greater turnover between years than we saw in the host between 2005-2007 and 2013-2018, however, between 2007-2013, turnover in the viruses is at a similar near-zero level as seen in GSB-A (Figure 3C). In GSB-B, we were only able to calculate turnover between 2005-2007 and 2007-2018 due to lack of coverage in the other years, but GSB-B had higher turnover compared to GSB-A (Figure 3C). Notably, the higher levels of SNP turnover in CV-1-33 were not restricted to the “variable” region. CV-1-33 is thus a uniquely dynamic genome, both at the gene content level (gene replacement from year-to-year) and SNP level (allele turnover in conserved genes) (Figure 3C).

### Theoretical model of contrasting strategies for GSB virus in Trout Bog Lake

Although GSB-A and GSB-B exist under the same ecological conditions with similar genomes (Table S3), these two hosts appear to experience different types of viral infections. GSB-A is associated with persistent prophages with high prevalence rate (infecting nearly 100% of the host population), while GSB-B is associated with two different viral types: a rare prophage infecting a subset of the population, and a (likely) lytic virus with a rapidly evolving genome. Because GSB-A went through a genome-wide sweep, which resulted in decreased population diversity, next we considered whether host population microdiversity could explain this difference in infection types, as it could impact the resistance potential for each host population. GSB-A had lower population diversity in most years examined, and we found that the most abundant viruses were prophages infecting the entire population. In scenarios like this with (nearly) clonal host populations, unless the fitness cost associated with each prophage is high, the probability of a resistant strain appearing will be low because there are fewer uninfected hosts with resistance potential existing in the population. From the virus perspective, the most successful strategy once most of the host population has been infected, is a long latent/lysogenic cycle that would allow a stable coexistence of virus and host. In contrast, a primarily lytic virus would likely lyse a substantial portion of their hosts, and a resistant host would eventually arise despite the host population being originally clonal.

GSB-B had higher population diversity in the years examined, so there is likely an existing pool of potentially resistant hosts that could be selected for. For this population, instead of stable prophages, we observe either a “rare” prophage, or a rapidly changing lytic virus that would be consistent with ongoing kill-the-winner/red-queen type dynamics. Hence, in higher-diversity host populations, the most successful strategy for viruses could be lytic or short lysogenic infection cycles, leading to more replication events and a more diverse viral population. Lysogenic viruses infecting these higher-diversity host populations, while less successful (in terms of relative abundance and infection rate), could still be stably associated with a subset of the host population as observed here in the GSB-B virus, CV-1-4.2.

To examine the potential for these scenarios to occur in nature, we adapted an existing theoretical model (Weitz et al., 2019) to consider the relationship between lysogeny, induction rate, and host resistance/susceptibility. As a first step, we were interested in identifying conditions that would allow for maximum lysogeny (majority of the host population are lysogenized), versus those favoring lytic activity (see Methods). While our model does not include a direct measurement for diversity, we used the initial abundance of lysogenized host cells as a simple proxy. Indeed, the genome-wide sweep in GSB-A was associated with an increase in the percentage of lysogenized cells; while we can’t distinguish if one directly caused the other, a population bottleneck event could lead to a higher ratio of lysogenized vs susceptible cells. Meanwhile, our model allows for hosts to transition from susceptible to resistant, but not from lysogenized to resistant, so resistance would have a greater chance to arise in a diverse population with a lower initial proportion of lysogenized hosts because there is a greater proportion of hosts that could become resistant.

Across the full landscape of lysis/induction rate and initial lysogen ratio, our model broadly recapitulated known virus-host dynamics. This model assesses the relationship between the initial lysogeny ratio (proportion of host population that is already lysogenized) and induction rate, and how those variables affect factors such as infection rate (lysogeny, specifically) and resistant host density. When a sufficient proportion of the initial host population (>20-40%) is already lysogenized at the start of the scenario, the model predicts higher rates of lysogenic infection compared to lower initial lysogeny ratios (Figure 4A, Figure S7). However, this infection rate is dependent on induction rate. If the induction rate drops too low, then lysogenic infection rate decreases due to prophage decay. On the other end, the higher infection rate with higher initial lysogeny ratio holds until the induction rate increases past a certain threshold. At this point, lysogenic infection rate drops, and resistant host density increases (Figure 4A, Figure S7). This suggests that there is a tipping point in this model, where a sufficiently high induction rate will allow the virus-host dynamics to enter into predator-prey/red-queen dynamics. This may create sufficient selection pressure from the viruses on the hosts, leading to an increase in the density of resistant hosts. When the induction rate is in an optimal medium range, then selection pressure for resistance on the hosts remains low, but induction is frequent enough to offset prophage decay.

**Figure 4.**
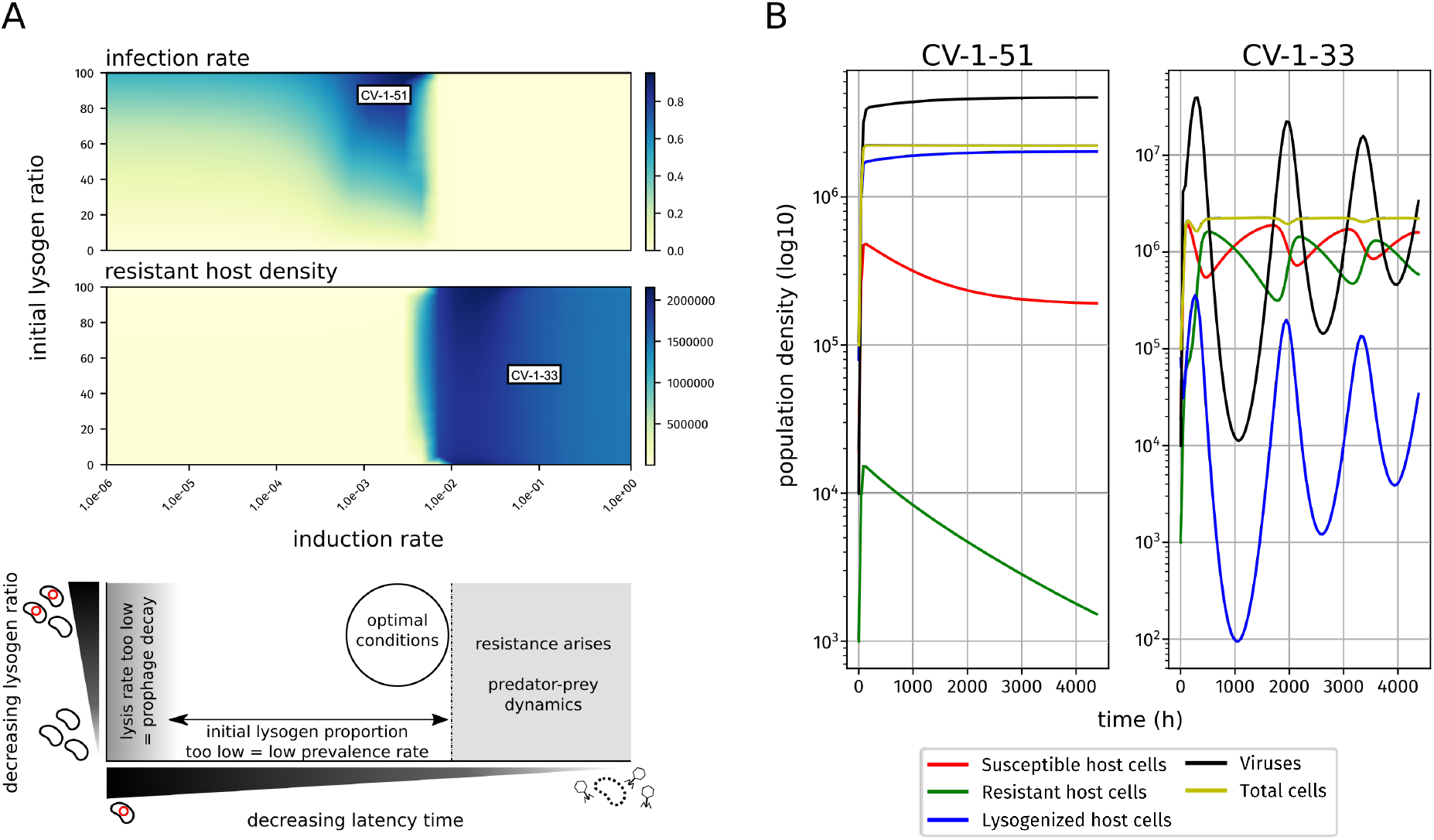
lysis rate against ratio of lysogenized host cells, showing infection pate (top) or resistant host density (center); color scales on the right for each respective plot; (bottom) shows a summary schematic for the above plots. CV-1-51 and CV-1-33 are labeled to represent the conditions that generated the vignettes. **(B)** vignettes for CV-1-51 (left) and CV-1-33 (right); population density for each respective group (see legend) against time (6 months total).

We can further examine these virus-host dynamics by modeling individual vignettes with specific conditions that (attempt to) mimic those from two of our GSB viruses, and deduce the possible scenarios that gave rise to the virus-host dynamics seen in our lake system. We first verified that our model recovered expected predator-prey dynamics for lytic viruses by testing a case similar to CV-1-33, a virus with low latency (i.e. mostly lytic) and low initial infection rate (Figure 4B). Here we see expected cyclical dynamics for viruses, resistant hosts, and susceptible hosts, consistent with the relative abundance patterns observed for CV-1-33 (Figure 4B).

Next, we used this model to investigate the conditions that may enable the long-term maintenance of a stable lysogen like CV-1-51. CV-1-51 infects most (if not all) of the GSB-A population, and has remained stable for over 10 years. In this scenario, we modeled a low induction rate, and started the time course with a high initial lysogeny/susceptible host ratio to mimic the conditions after the genome sweep, because most of the host population appears to be infected with CV-1-51. For the 6 months we modeled, we find a slow, but steady increase in lysogenized host cells, and no significant rise of resistant host cells (Figure 4B). Hence, with low levels of induction, and following a genome-wide sweep which would have increased its prevalence, CV-1-51 can be maintained within a near-clonal population without viral genome decay nor triggering a rise in host resistance. While more sophisticated mechanisms for this type of stability may exist, such as density dependence induction rate (Stokar-Avihail et al., 2019), we did not find evidence of those mechanisms occurring here, and our model suggests that this type of stability is possible even without those.

## CONCLUSION

In this study, we characterize viruses infecting Green sulfur bacteria (GSB) in a model freshwater lake. By leveraging the unique autofluorescence of GSB cells, we sorted GSB cells using FACS, and sequenced these “targeted metagenomes” to identify GSB-specific viruses. These targeted metagenomes allowed us to observe “in-cell” viruses, and ecologically relevant ongoing infections. Overall, this newly developed method recovered novel viruses with a host context at high (population-level) resolution. These new host-contextualized viruses in turn enabled a reanalysis of time series bulk metagenomes to investigate virus-host interactions across seasons and years in a model freshwater lake. Notably, we observed how a GSB host population stably co-existed with two distinct prophages that reached a consistent ~100% infection rate over multiple years without seemingly triggering resistance in the host population. Our data further suggest that host population diversity may be a critical factor in driving virus-host interactions, by influencing the potential for resistance in the host population. Consequently, the optimal latency time decision of viruses would in turn depend partly on the host population diversity, with low-diversity host populations more closely linked to long-latency lysogens. This would be consistent with the link between lysogeny and host life-history traits, as lysogeny-associated traits such as “pathogenicity” or “boom-and-bust” life cycles are also associated with population bottlenecks, and could also explain the imperfect linkage between lysis-lysogeny rate and ecological parameters such as host abundance and growth rate.

## ACKNOWLEDGEMENTS

We thank Nandita Nath for her work generating the sequencing libraries, Sean Jungbluth for help with genome binning for our 2017 pilot data, and Charles Olmsted and Kali Denis for their work in the field to collect the lake samples. We also thank Donald Bryant and Jennifer Thweatt from the Bryant Lab for kindly providing Chlorobi isolates for FACS benchmarking and sharing their knowledge of these bacteria. This work was conducted by the US Department of Energy Joint Genome Institute, a Department of Energy Office of Science User Facility under contract no. DE-AC02-05CH11231, and used resources of the National Energy Research Scientific Computing Center, which is supported by the Office of Science of the US Department of Energy under contract no. DE-AC02-05CH11231. This work was supported by the Laboratory Directed Research and Development Program of Lawrence Berkeley National Laboratory under US Department of Energy contract no. DE-AC02-05CH11231.

