## Supplementary Tables (2-4) and Figures for "Host population diversity as a driver of viral infection cycle in wild populations of green sulfur bacteria with long standing virus-host interactions"

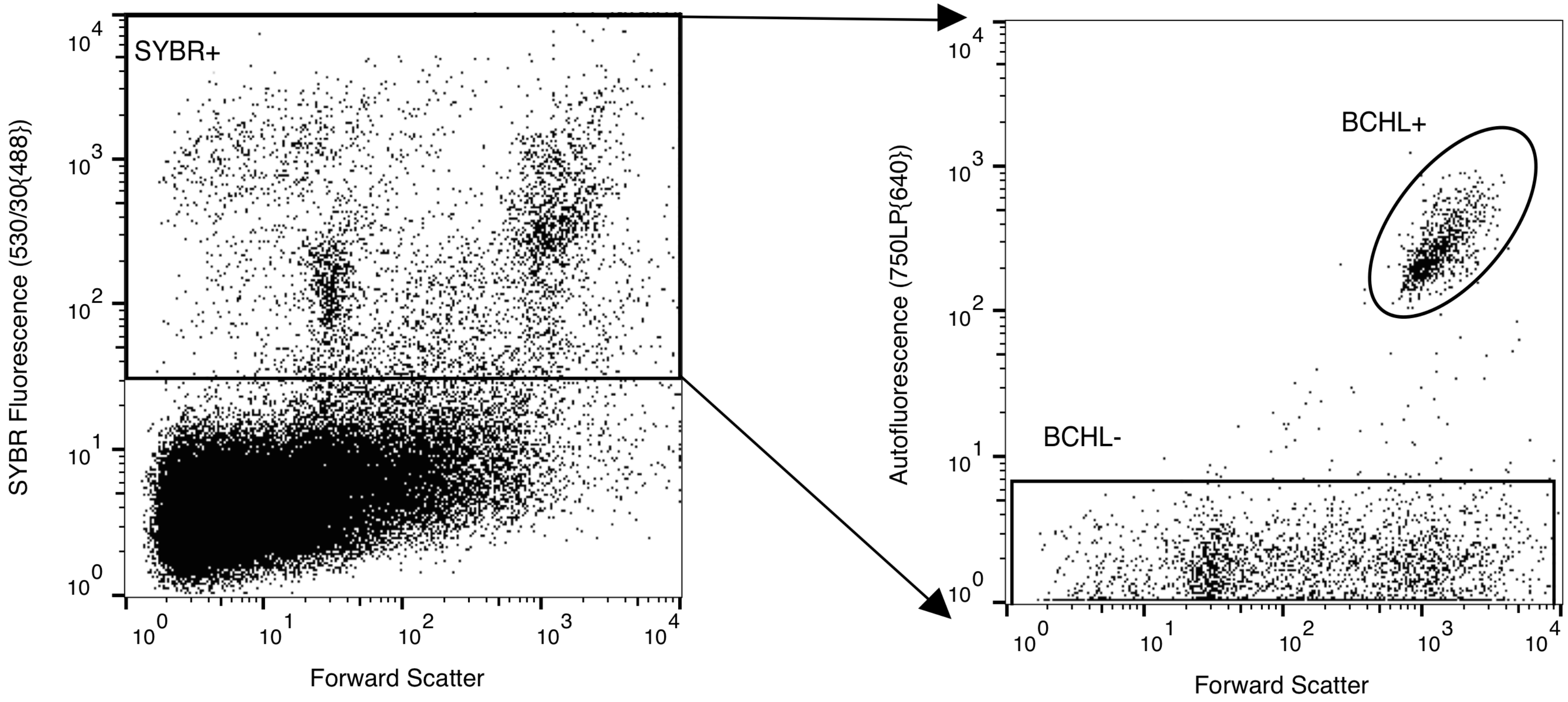

**Figure S1.** The total cell population of samples were identified by staining with SYBR Green II, and gating off 530/40 fluorescence when excited by a 488nm laser. Within this subset of events, bacteriochlorophyll positive and negative populations were distinguished by the presence or absence of far-red (750LP) autofluorescence when excited by a 640nm laser. Pictured above is a representative image from the 2.5 meter sample harvested on Aug 13, 2018. All samples contained at least one decade of separation on the 750LP channel between the BCHL+ and – gates.

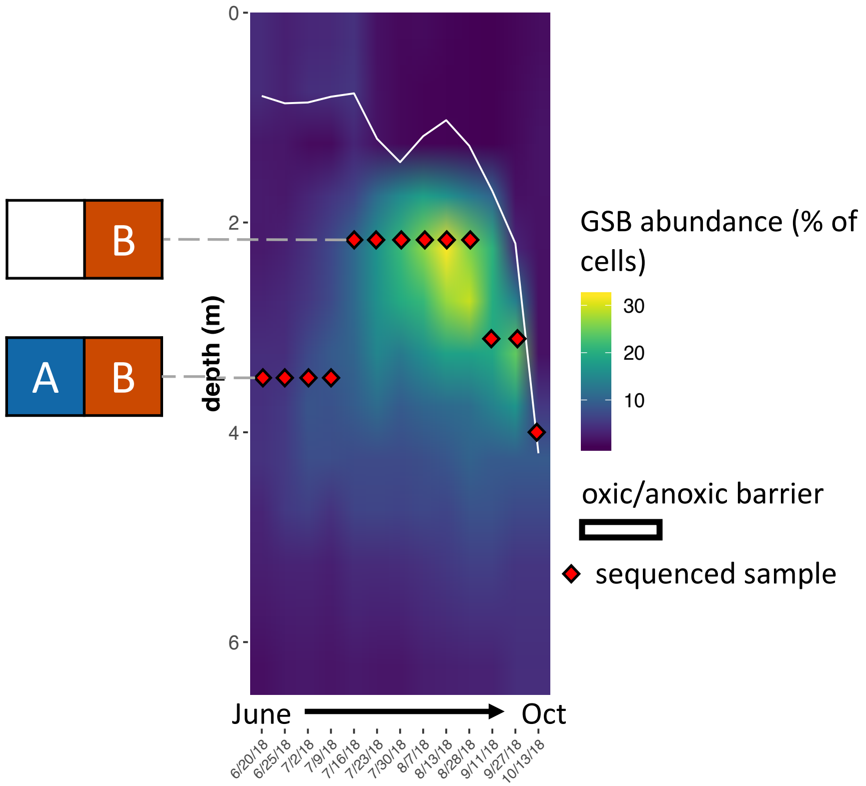

**Figure S2.** An average of 3.7 million cells/mL were measured for per sample per date; shown is GSB abundance (% of total cells per sample per date). Red diamonds represent the specific depth that was sequenced. The white line represents the oxic/anoxic barrier. GSB-A (A) was only present in samples below 3m, while GSB-B (B) was present at all depths sequenced.

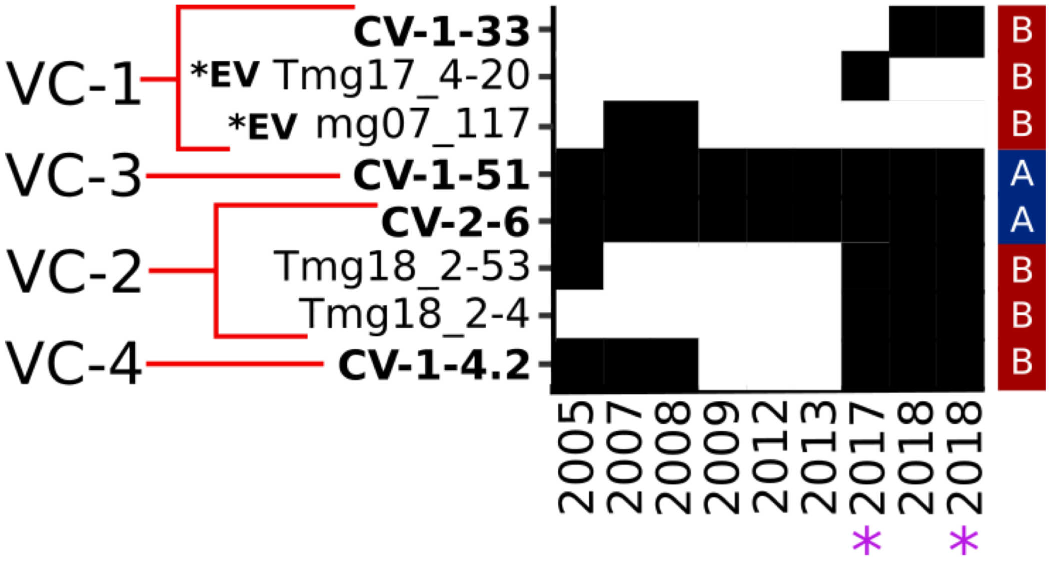

**Figure S3.** Shown are presence/absence values for viral clusters shown in red in Figure 2. The four viral contigs discussed in this paper are labeled in bold; variants of CV-1-33 are labeled as *EV in bold. Shown is presence/absence for each year across all contigs; starred years represent the targeted metagenomes, where at least half of all replicates must contain the contig to be counted as present that year. Hosts are labeled on the right.

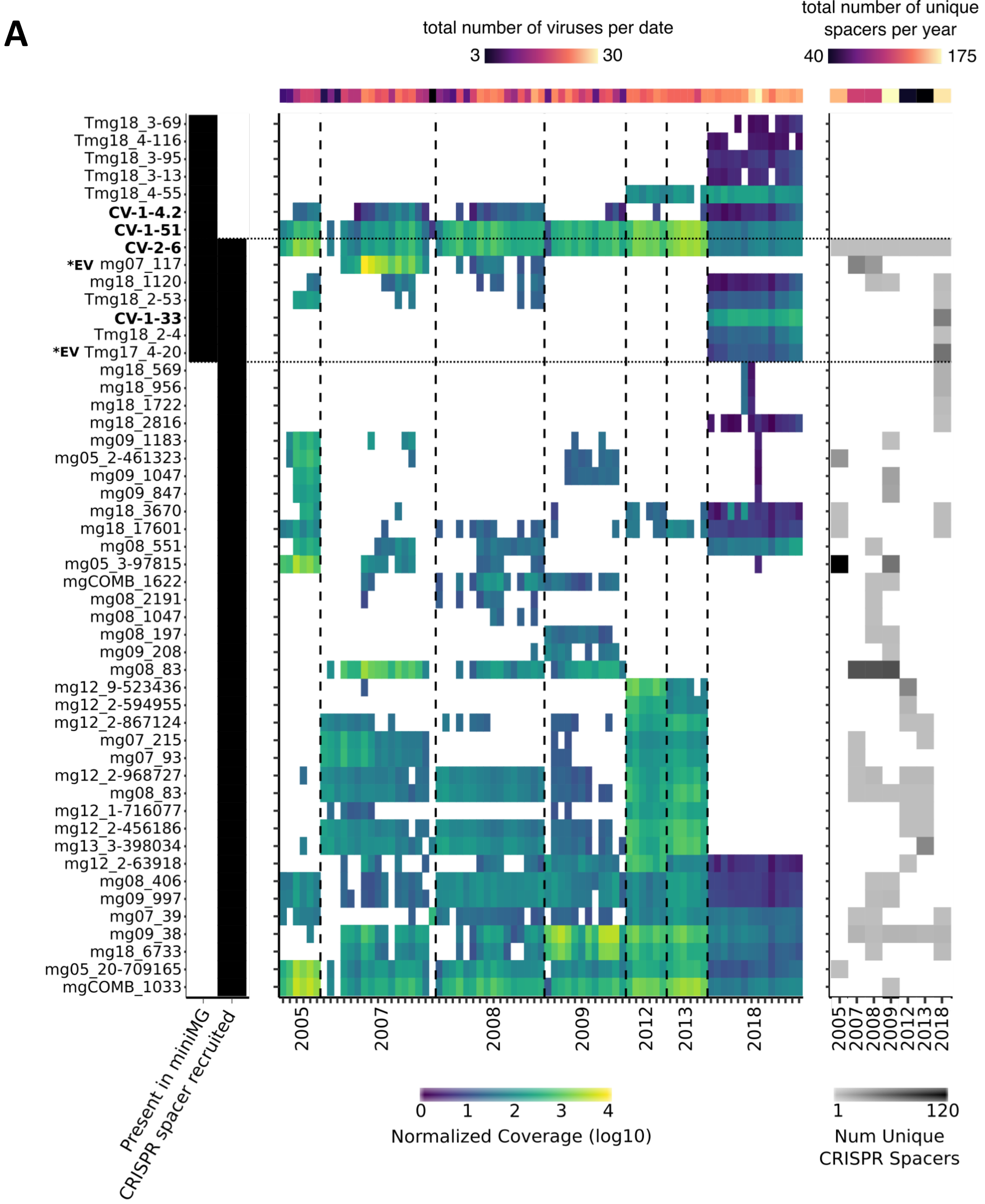

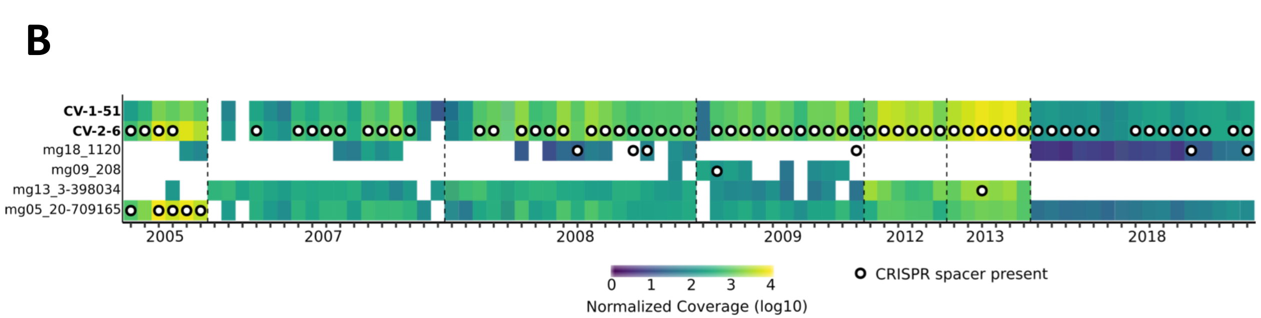

**Figure S4. (A)** Shown are viral contigs identified through GSB CRISPR spacer matching, plus the identified GSB viral contigs through FACS flow sorting. The four viral contigs discussed in this paper are labeled in bold; variants of CV-1-33 are labeled as *EV in bold. *(left)* Black to signify which contigs were found in the targeted metagenomes (miniMG), and which contigs were recruited through CRISPR spacer matching. *(center)* Normalized coverage for each sample across all contigs; white space represents insufficient or no coverage for that contig/sample. *(right)* Shown are the number of unique CRISPR spacers acquired for each year. **(B)** Shown are the same coverage values for GSB viral contigs, plus white with black circles that represent the presence of a CRISPR spacer.

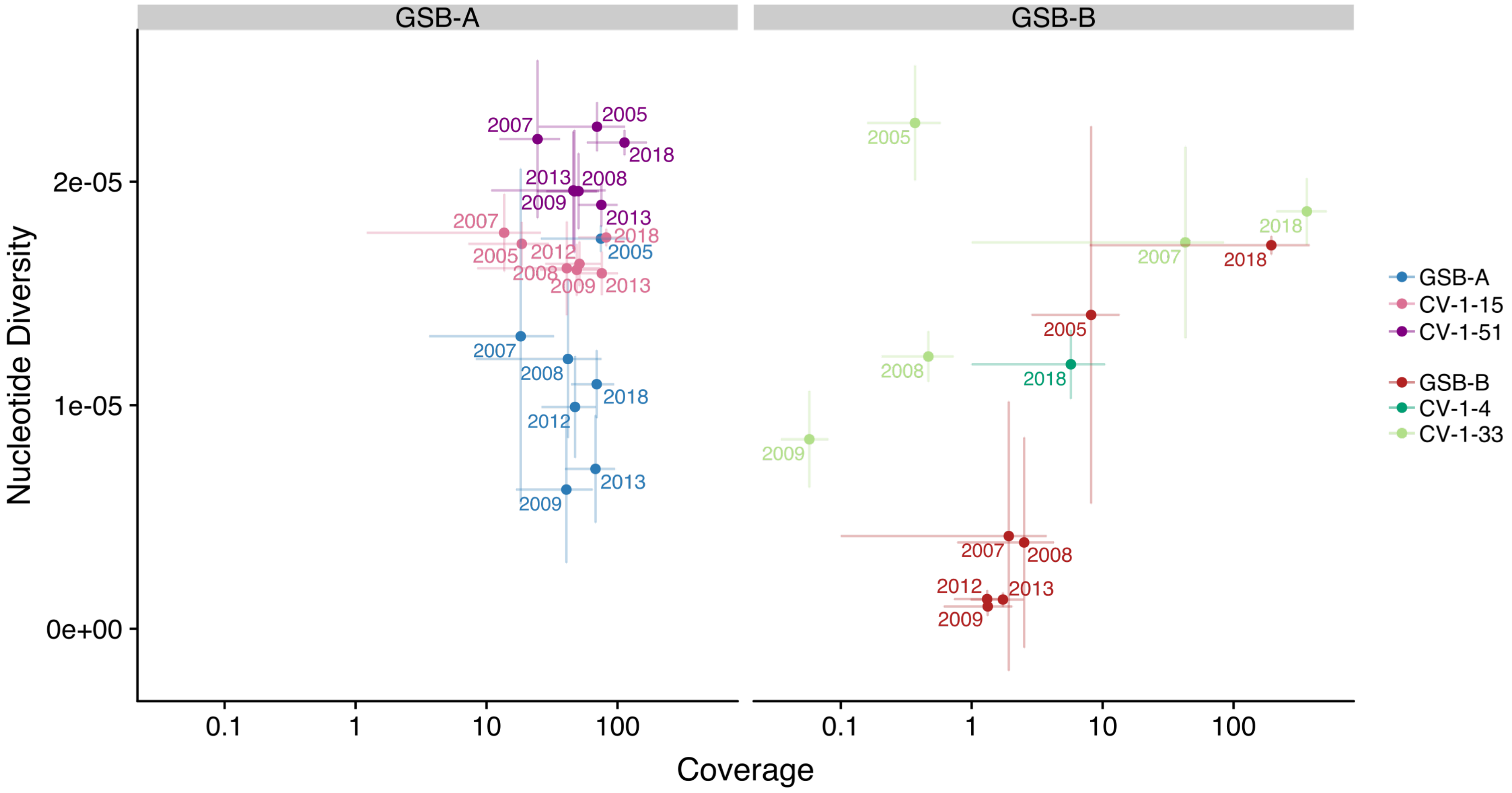

**Figure S5.** Nucleotide diversity of GSB-A, GSB-B, and their associated viruses against coverage (log-scale; not normalized); bars show standard deviation.

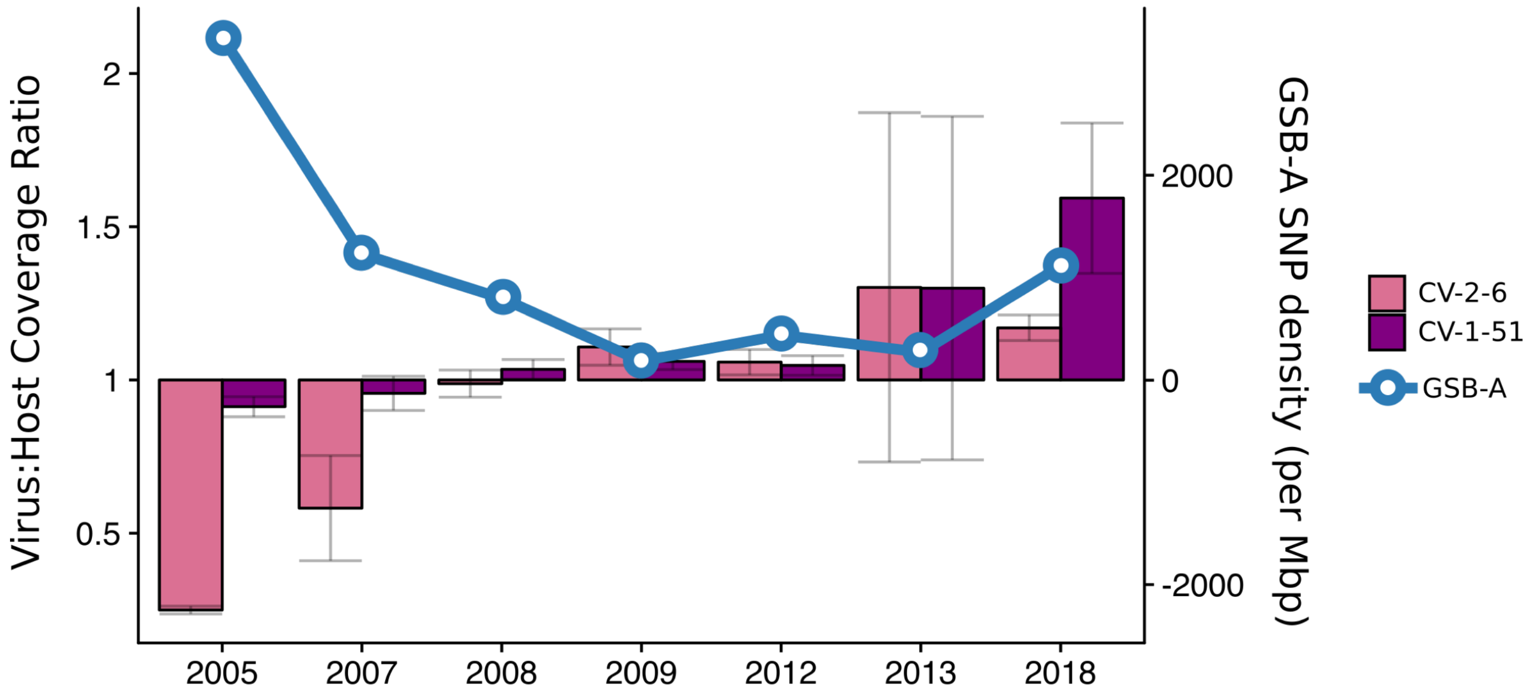

**Figure S6.** Shown are ratio of virus:host normalized coverage values of CV-2-6 and CV-1-51; error bars are standard deviation. Line represents the SNP density (SNPs per Mbp) for GSB-A.

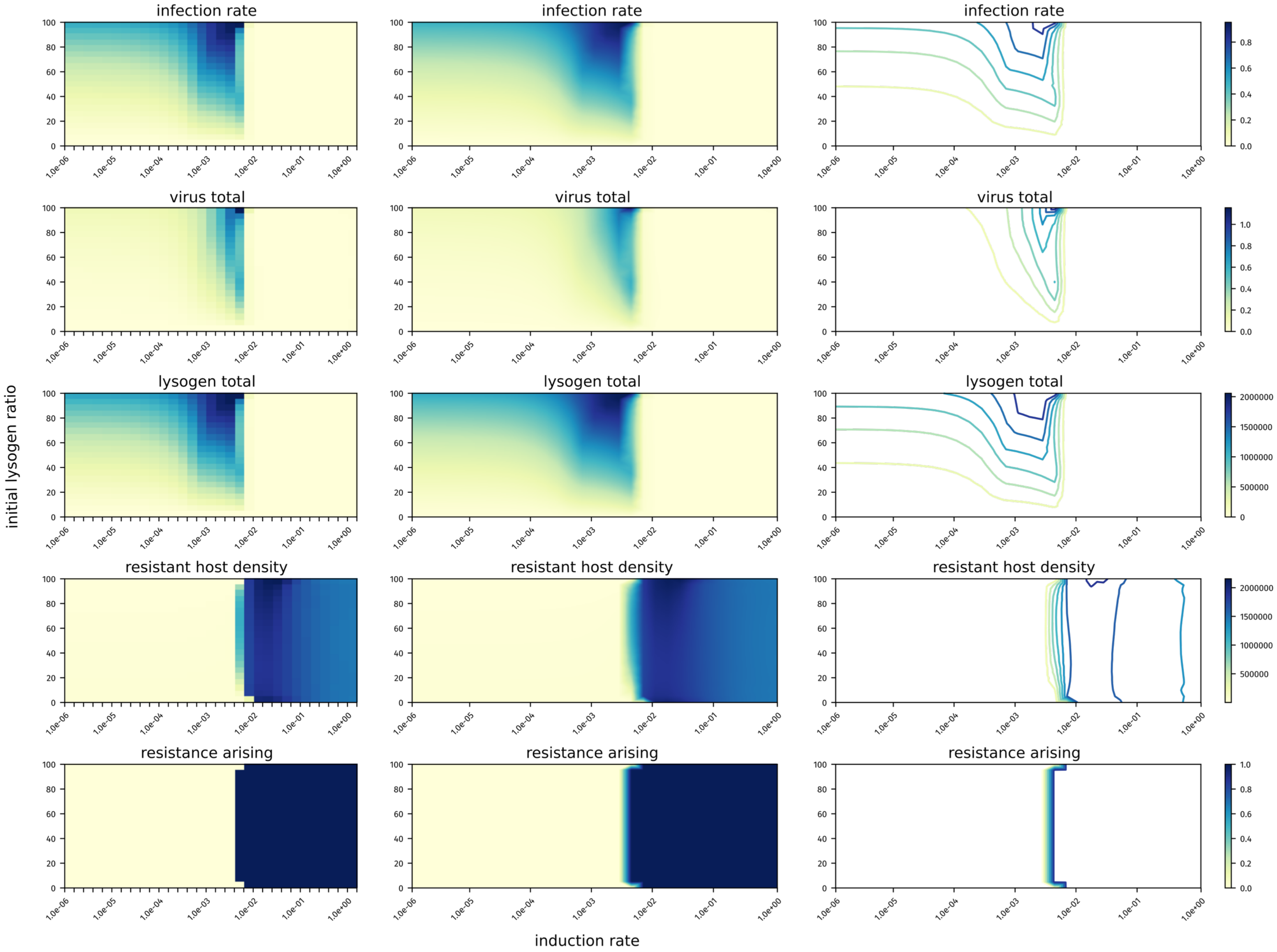

**Figure S7.** Induction rate against ratio of lysogenized host cells, showing infection rate (top), virus total (virion plus lysogens), lysogeny total, resistant host density, resistance arising (bottom); color scales on the right for each respective plot; shown raw data (left column), interpolation (center), and interpolation contour (right).

| | GSB-A | GSB-B | Sampling Depth (m) |
|---|---|---|---|
| 11-Jun | (98.9/2.75) | (99.04/0.55) | 3.5 |
| 20-Jun | (98.9/1.1) | (99.45/1.65) | 3.5 |
| 25-Jun | (99.45/2.2) | (98.9/1.1) | 3.5 |
| 2-Jul | (99.45/3.02) | (98.08/2.2) | 3.5 |
| 9-Jul | (98.9/1.1) | (100/2.2) | 3.5 |
| 16-Jul | | (99.45/0.55) | 2 |
| 23-Jul | | (99.45/1.65) | 2 |
| 30-Jul | | (98.35/1.92) | 2 |
| 7-Aug | | (99.45/1.1) | 2 |
| 13-Aug | | (99.45/1.65) | 2 |
| 20-Aug | | (99.04/0.55) | 2 |
| 11-Sep | | (98.9/1.65) | 3 |
| 27-Sep | (35.16/0) | (98.9/1.65) | 3 |
| 13-Oct | (98.76/1.1) | (99.31/2.2) | 4 |

**Table S2.** Genome bins assembled for each sampling date. Shown are (Completion / Redundancy) for each bin, along with the sampling depth for the sampling date.

**A**

| | Completeness | Redundancy |
|---|---|---|
| 2018 GSB-A | 99% | 2% |
| 2018 GSB-B | 99% | 2% |

**B**

| ANI | 2005-13 GSB-A (Chlorobium-111) | 2005-13 GSB-B (Chlorobium-3520) | 2017 GSB-A | 2017 GSB-B |
|---|---|---|---|---|
| 2018 GSB-A | **0.995985069** | 0.849017794 | **0.997858344** | 0.851399955 |
| 2018 GSB-B | 0.850665765 | **0.998608488** | 0.854558088 | **0.999801978** |

**C**

| AF | 2005-13 GSB-A (Chlorobium-111) | 2005-13 GSB-B (Chlorobium-3520) | 2017 GSB-A | 2017 GSB-B |
|---|---|---|---|---|
| 2018 GSB-A | **0.737023826** | 0.431735066 | **0.987421399** | 0.506435925 |
| 2018 GSB-B | 0.460059715 | **0.752216605** | 0.454740141 | **0.938266211** |

**Table S3.** Information regarding the 2 genome bins identified in 2018 from Trout Bog Lake; **A)** genome completeness and redundancy, **B)** ANI compared to previously identified genome bins from Trout Bog Lake, **C)** alignment fraction (AF, percent) compared to previously identified genome bins from Trout Bog Lake

| Contig Name | length (bp) | Predicted Host |
|---|---|---|
| CV-1-51 | 14619 | GSB-A |
| CV-2-6 | 41723 | GSB-A |
| CV-1-33 | 33736 | GSB-B |
| CV-1-4.2 | 31527 | GSB-B |
| Tmg18_4-55 | 41317 | GSB-B |
| Tmg18_4-116 | 37298 | GSB-B |
| Tmg18_3-013 | 11633 | GSB-B |
| Tmg18_2-4 | 29817 | GSB-B |
| Tmg18_2-53 | 28582 | GSB-B |
| Tmg18_3-69 | 11643 | GSB-B |
| Tmg18_3-94 | 12828 | GSB-A |
| Tmg18_3-84 | 15194 | GSB-B |
| Tmg18_2-59 | 8364 | GSB-A |
| Tmg18_4-56 | 42242 | GSB-B |
| Tmg18_4-73 | 14738 | GSB-B |
| Tmg18_4-94 | 5072 | GSB-B |
| Tmg18_4-124 | 3820 | GSB-B |
| Tmg18_4-133 | 36543 | GSB-B |
| Tmg18_4-148 | 10084 | GSB-B |
| Tmg18_4-176 | 9431 | GSB-B |
| Tmg18_4-219 | 2800 | GSB-B |
| Tmg18_1-99 | 13477 | GSB-B |
| Tmg18_3-33 | 6581 | GSB-B |
| Tmg18_3-45 | 8091 | GSB-B |
| Tmg18_3-9 | 7105 | GSB-B |
| Tmg18_3-18 | 6854 | GSB-B |
| Tmg18_3-36 | 5173 | GSB-B |
| Tmg18_1-35 | 8168 | GSB-B |
| Tmg18_2-88 | 12203 | GSB-B |
| Tmg18_2-107 | 5140 | GSB-B |
| Tmg18_2-125 | 15336 | GSB-B |
| Tmg18_2-029 | 8972 | GSB-B |
| Tmg18_2-077 | 7513 | GSB-B |
| Tmg18_3-6 | 9941 | GSB-B |
| Tmg18_2-1 | 26472 | GSB-B |
| Tmg18_2-10 | 6186 | |
| Tmg18_2-26 | 7679 | |
| Tmg18_3-38 | 3113 | GSB-B |
| Tmg18_3-82 | 8648 | GSB-B |
| Tmg18_3-117 | 5759 | GSB-B |
| Tmg18_3-121 | 10364 | GSB-B |
| Tmg18_3-47 | 5823 | GSB-A |
| Tmg18_3-81 | 3238 | GSB-A |

**Table S4.** Shown are non-redundant contigs predicted to be viral contigs using VirSorter. The first block contains the 11 viral contigs that were identified trhough manual curation as complete or near-complete. The bottom block are putative viral contigs that were considered too short or too decayed to validate as viral genomes. Contigs with GSB-A as their predicted host are shaded in grey for convenience.
